# Subcellular Localization of Dopamine D1 and D2 Receptors in the Mouse Hippocampus

**DOI:** 10.64898/2026.04.23.720385

**Authors:** Cameron Swope, Garrett Sommer, Ravon Smith, Teresa A. Milner, Jimcy Platholi

**Author notes:** Corresponding author for journal: Cameron Swope 1300 York Ave, New York, NY 10065. Corresponding authors for manuscript: Jimcy Platholi, 1300 York Ave, New York, NY 10065, Teresa A. Milner, Feil Family Brain and Mind Research Institute, Weill Cornell Medicine, 407 East 61^st^ St room 307, New York, NY 10065.

## Abstract

Dopamine signaling through dopamine 1 receptors (D1R) and dopamine 2 receptors (D2R) regulates hippocampal synaptic plasticity underlying learning and memory, yet their subcellular localization within the hippocampus is unknown. Here we performed electron microscopic immunocytochemistry to elucidate the distribution of D1R and D2R in subregions of the mouse hippocampus. In CA1 and CA3 stratum radiatum (SR), D1R- and D2R-immunoreactivity was found primarily on pyramidal cell dendritic spines and unmyelinated axons, and to a lesser extent in axon terminals and glia. In both regions, D1R-labeled terminals formed predominantly asymmetric (excitatory-type) synapses on dendritic spines, whereas D2R-labeled terminals formed mainly symmetric (inhibitory-type) synapses on pyramidal cell dendritic shafts. In the dentate gyrus (DG) hilus, D1R-labeling was almost exclusively found in unmyelinated axons and glia. D2R immunoreactivity in the hilus similarly was present in unmyelinated axons and glia but was also detected in dendritic spines originating from mossy cells and in terminals forming symmetric synapses. These findings indicate that dopamine receptors are positioned to influence excitatory and inhibitory signaling in the murine hippocampus. As D1R and D2R exert opposing effects on neuronal signaling, their localization on pyramidal neuron compartments provides a structural substrate for bidirectional modulation of synaptic plasticity and pyramidal cell activity. In addition, the presence of D2Rs on inhibitory terminals contacting pyramidal neurons and hilar interneurons suggests a role in regulating inhibitory circuitry within the hippocampus.

## Introduction

Dopamine is a neuromodulator that regulates neuronal excitability, synaptic plasticity, and network dynamics across the central nervous system (CNS)^[1, 2]^. Dopaminergic signaling exerts its effects primarily through dopamine receptors (DARs), a family of G-protein coupled receptors (GPCRs) that modulate intracellular signaling cascades and neurotransmitter release^[3, 4]^. Five DARs are expressed in the CNS and are broadly classified into two functional families: D1-like receptors (D1, D5), which stimulate adenylyl cyclase, and D2-like receptors (D2, D3, D4) which inhibit adenylyl cyclase activity^[5]^. Through these signaling pathways, DAR activation bidirectionally regulates neuronal firing, synaptic transmission, and plasticity^[3]^, and dysregulation of DAR signaling represents a core mechanism underlying multiple neurodegenerative and neuropsychiatric disorders^[6, 7]^.

DAR-mediated modulation of neuronal function depends not only on receptor subtype but also on cell-type-specific expression and subcellular localization. DARs are expressed by both glutamatergic neurons and diverse classes of GABAergic interneurons, where they localize to distinct presynaptic, axonal, and postsynaptic compartments^[8–15]^. Consequently, the functional impact of dopaminergic signaling within neural circuits is determined by the subcellular distribution of DARs. Axonal DARs regulate action potential propagation^[15]^, presynaptic DARs control neurotransmitter release probability and vesicle dynamics^[8, 9]^, and postsynaptic DARs modulate dendritic integration^[16]^, intrinsic excitability^[17, 18]^, and synaptic plasticity^[19]^.

In the hippocampus, dopamine modulates synaptic strength and is a critical regulator of synaptic plasticity underlying learning and memory^[20–24]^. Dopaminergic signaling gates both long-term potentiation (LTP) and long-term depression (LTD) by regulating postsynaptic depolarization^[18]^, NMDA and AMPA receptor function^[11]^, and downstream calcium-dependent signaling cascades essential for plasticity induction and stabilization^[25, 26]^. Through these mechanisms, dopamine promotes activity-dependent recruitment of neurotrophic signaling required for memory consolidation and circuit remodeling^[27]^. Consistent with these roles, hippocampal dopaminergic signaling is required for spatial and contextual learning^[28]^, memory persistence, and cognitive flexibility^[29, 30]^, and contributes to maladaptive plasticity associated with reward learning, addiction-related associative memory formation and reinstatement^[31, 32]^, as well as neuropsychiatric and neurodegenerative disorders. Importantly, dopamine’s effects within the hippocampus are highly heterogenous, reflecting synapse and subcellular distribution-specific differences in DAR subtype expression across hippocampal subregions and neuronal populations.

Studies based on reporter mice suggest that DAR expression in the hippocampus is region- and cell-type-specific. In CA1 and CA3, pyramidal neurons may preferentially express D1 receptors (D1R), while interneurons express both D1 and D2 receptors (D2R) in layer-specific patterns^[33–35]^. In the dentate gyrus (DG), granule cells and GABAergic hilar interneurons express D1Rs, whereas D2R expression is enriched in mossy cells^[34]^. Although these studies provide important insights, they do not establish the subcellular localization of endogenous DAR proteins, an essential determinant of receptor function. Thus, despite extensive evidence that dopamine critically shapes hippocampal plasticity and behavior, a comprehensive, ultrastructural understanding of DAR subtype localization across hippocampal circuits is lacking. This gap limits our ability to predict how dopamine release modulates synaptic transmission and plasticity within specific hippocampal subregions and cell types.

To address this, we used electron microscopic immunohistochemistry to quantify the subcellular distribution of D1R and D2R in the CA1, CA3, and DG subregions in male and female mice. By assessing the region-, cell-type-, synaptic- and sex-specific differences of D1R and D2R, this study provides an expansive framework for understanding how dopamine signaling is organized at the synaptic level in the dorsal hippocampus.

## Materials and Methods

### Animals

Experimental procedures were approved by the Institutional Animal Care and Use Committee of Weill Cornell Medicine and were conducted in accordance with the National Institute of Health Guide for the Care and Use of Laboratory Animals (8^th^ edition, 2011). A total of six 2-month-old C57BL/6J mice (3 males and 3 females) were used for these studies. Mice were housed 3-4 per cage on a 12-hour light/dark cycle with *ad libitum* access to food and water. Animals were allowed to acclimate to the vivarium prior to euthanasia. The tissue used in this study were derived from brains collected from a previous cohort described previously (Marques-Lopes et al. JCN 522:3075, 2014).

### Tissue Preparation

Tissue was prepared using previously described procedures^[36]^. Mice were deeply anesthetized with sodium pentobarbital (150 mg /kg, i.p.) and transcardially perfused with 0.9% saline containing 2% heparin, followed by 30 mL of fixative consisting of 3.75% acrolein (Polysciences, Washington, PA) and 2% paraformaldehyde (PFA; Electron Microscopy Sciences (EMS), Fort Washington, PA, cat # 19208) in 0.1M phosphate buffer (PB), pH 7.4. Brains were removed and post-fixed for 30 minutes at room temperature on a shaker in a solution containing 1.9% acrolein and 2% PFA in PB. Brains were coronally blocked between the caudal hippocampus and pons using a brain mold (Activational Systems Inc., Warren, MI). Coronal sections (40 μm thick) were cut on a VT1000X vibratome (Leica Microsystems, Buffalo Grove, IL) and stored at −20 ◦C in cryoprotectant solution (30% sucrose, 30% ethylene glycol in PB) until immunocytochemical processing.

### Antibody Characterization

#### D1R

A rat monoclonal antibody against dopamine D1 receptor (D2944, Sigma-Aldrich, RRID: AB_1840787) was used at a 1:750 dilution (**Table 1**). Antibody specificity has been previously validated by 1) absence of Western blot signal at ∼95 and ∼100 kD in brain homogenates from *drd1* (-/-) knockout mice, 2) absence of immunofluorescence labeling in tissue from *drd1* (-/-) knockout mice, and 3) immunoprecipitation of D1R, but not D2R, protein confirmed by mass spectrometry^[37]^. This antibody has been previously published in immunohistology studies^[38]^ including prior electron microscopy studies^[39]^.

**Table 1:**
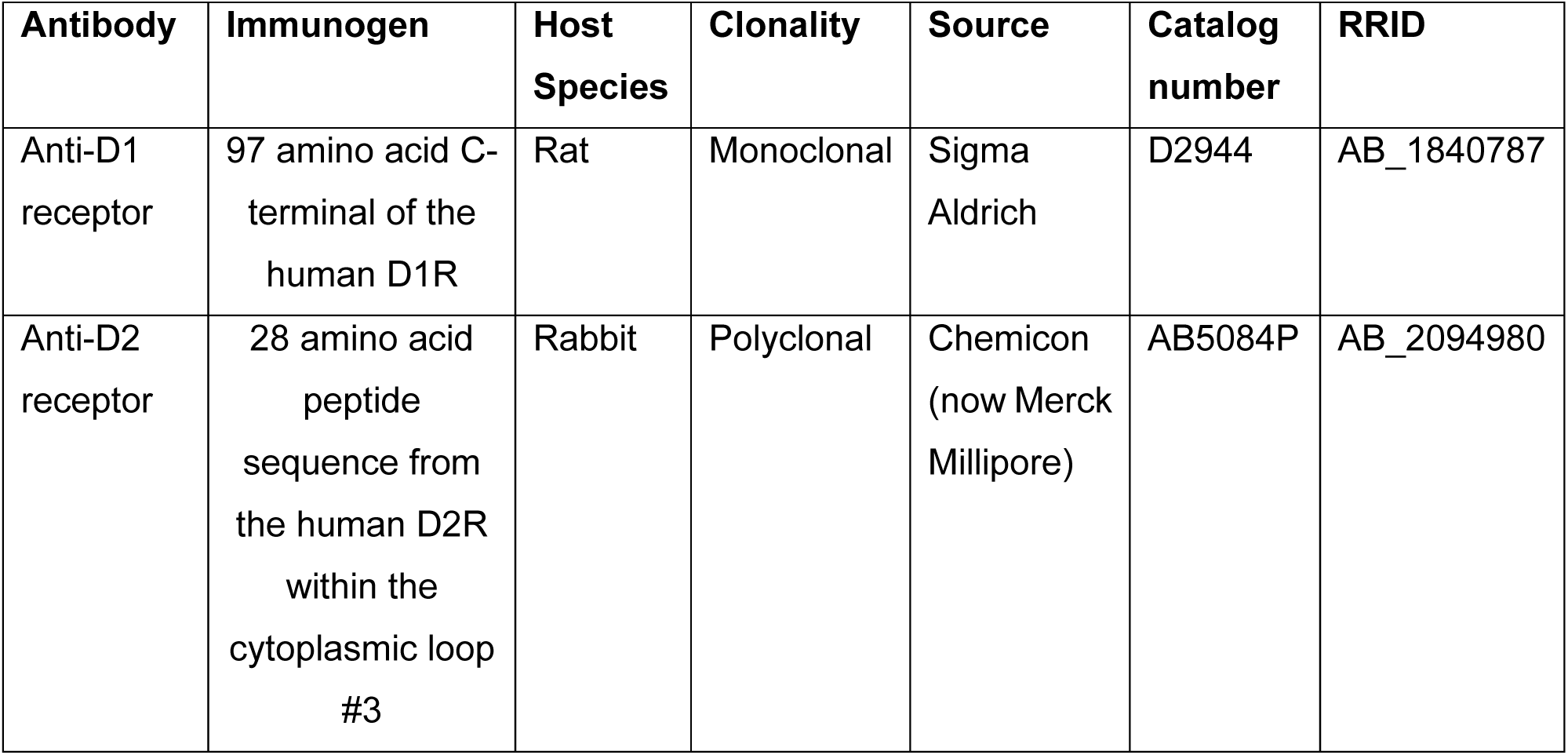
Antibody Information.

| Antibody | Immunogen | Host Species | Clonality | Source | Catalog number | RRID |
| --- | --- | --- | --- | --- | --- | --- |
| Anti-D1 receptor | 97 amino acid C-terminal of the human D1R | Rat | Monoclonal | Sigma Aldrich | D2944 | AB_1840787 |
| Anti-D2 receptor | 28 amino acid peptide sequence from the human D2R within the cytoplasmic loop #3 | Rabbit | Polyclonal | Chemicon (now Merck Millipore) | AB5084P | AB_2094980 |

#### D2R

A rabbit polyclonal antibody against dopamine D2 receptor (AB5084P, Chemicon (now Merck Millipore), RRID: AB_2094980) was used at a 1:300 dilution (**Table 1**). Specificity has been demonstrated previously by 1) absence of Western blot signal between ∼50 and ∼100kD in brain homogenates from *drd2* (-/-) knockout mice, 2) absence of immunofluorescence labeling in tissue from *drd2* (-/-) knockout mice, and 3) immunoprecipitation of D2R, but not D1R, protein confirmed by mass spectrometry^[37]^. This antibody has been previously published in immunohistology studies^[40]^ including prior electron microscopy studies^[41]^.

### Electron Microscopy Immunocytochemistry

Dorsal hippocampal sections were processed for electron microscopy using previously described protocols^[36]^. To ensure identical labeling between sexes, tissue sections were coded with hole punches in the cortex and pooled into common containers so that all samples were processed together through each immunocytochemical procedure. Sections were rinsed in PB and treated with 1% sodium borohydride in PB for 30 minutes to reduce residual aldehydes. Sections were then washed extensively in PB until no gas bubbles remained and incubated sequentially in: 1) 0.5% bovine serum albumin (BSA) in Tris-buffered saline (TS) for blocking; 2) primary antibody (D1R, 1:750; D2R, 1:300) diluted in BSA for 24 hours at room temperature followed by 24 hours at 4 °C; 3) biotinylated secondary antibody (1:400; donkey anti-rat IgG (D1R) or goat anti-rabbit IgG (D2R); Jackson ImmunoResearch Inc., West Grove, PA) for 30 minutes; and 4) avidin-biotin complex (ABC) solution for 30 minutes at half the manufacturer’s recommended concentration (Vector Laboratories, Burlingame, CA). All incubations and washes were performed on a shaker at 145 RPM. Bound peroxidase was visualized using a timed reaction in 3,3′-diaminobenzidine (DAB; Sigma-Aldrich Chemical Co., Milwaukee, MI) and hydrogen peroxide in TS (9 minutes for D1R; 7.5 minutes for D2R). Sections were thoroughly rinsed in TS between each incubation.

Following immunocytochemistry, sections were post-fixed in 2% osmium tetroxide in PB for 1 hour, dehydrated through graded ethanol and propylene oxide, and incubated overnight in a 1:1 mixture of propylene oxide and EMbed 812 epoxy resin (Electron Microscopy Sciences, EMS). Sections were then incubated in pure Embed 812 for 2 hours and flat-embedded between sheets of Aclar plastic, followed by polymerization at 60°C for 3 days. Serial ultrathin sections (70 nm) through the region of interest (ROI) were cut using a diamond knife (EMS) on an ultramicrotome (Leica EM UCT6) and collected onto mesh thin-bar copper grids (T400-Cu, EMS). Sections were counterstained with Uranyless™ (catalog #22409, EMS) followed by lead citrate (Catalog #22410, EMS) to enhance ultrastructural contrast for electron microscopic examination.

### Imaging and Analysis

Experimenters were blinded to experimental groups during imaging and analysis. For each animal, ultrathin sections at the tissue-plastic interface were selected to minimize variability in antibody penetration. Electron micrographs were acquired using a Hitachi HT7800 transmission electron microscope (Hitachi High-Tech, Minato, Japan) equipped with a digital camera system (version 3.2, Advanced Microscopy Techniques, Woburn, MA). For quantification of DAR-labeled profiles, all immunolabeled profiles (e.g., dendritic spines, dendrites, axons, terminals, and glia) were photographed at 10,000-20,000x magnification within two grid squares (total area 6050 μm²) per mouse per hippocampal region. Three regions were sampled: 1) CA1 stratum radiatum (SR), 50-150 μm below the pyramidal cell layer; 2) CA3 SR, 50-150 μm above the pyramidal cell layer; and 3) the central hilus of the DG. Additionally, 50 randomly selected mossy fiber profiles from CA3 stratum lucidum (SLu) were imaged per animal to determine the proportion of mossy fibers containing DARs.

Immunolabeled profiles were classified according to established ultrastructural criteria^[42]^.

#### Dendrites

Dendritic profiles contained regular microtubule arrays and measured between 0.5-2.0 μm in cross sectional diameter. Pyramidal cell dendrites in CA1 and CA3 were often >1 μm in diameter, had dendritic spines, and received relatively few terminal contacts on the dendritic shafts. Interneuron dendrites in the DG usually had a diameter between 0.5 and 1 μm, lacked spines, and received multiple terminal contacts. Mossy cell dendrites in the DG hilus were generally > 1 μm in diameter and contained dendritic spines. Dendrites that did not meet these criteria above, were classified unidentifiable.

#### Dendritic spines

Dendritic spines were typically smaller than 0.2 μm in diameter, contacted by terminals forming asymmetric synapses identified by a robust postsynaptic density, and sometimes were contiguous with dendritic shafts.

#### Terminals

Axon terminals contained numerous small synaptic vesicles, mitochondria, and measured >0.2 μm in cross-sectional diameter. The term “contact” included asymmetric synapses, symmetric synapses and appositions. Asymmetric (excitatory-type) synapses formed by terminals were identified by clear postsynaptic densities on their targets. Symmetric (inhibitory-type) synapses formed by terminals were identified by thin pre- and post-synaptic densities. Appositions were defined as membrane contacts not separated by glial profiles but lacking intercleft material or conventional synapses in the plane of section analyzed.

#### Axons

Unmyelinated axons were profiles smaller than 0.2 μm in diameter, contained few synaptic vesicles, and had no apparent synaptic junction in the plane of section. *En passant* terminals had a clear synapse site with clustered synaptic vesicles, and unmyelinated axon protrusions on either side.

#### Glia

Glial profiles were identified by their tendency to conform to the shape of surrounding structures and/or the absence of microtubules.

#### Unknown

Profiles that could not be confidently identified were categorized as unknown.

### Figure preparation

Micrograph resolution was reduced to 300 dpi before sharpness, contrast, and brightness were adjusted in Image J. Image adjustments were made to the entire image without alterations to the original content of the raw images. Diagrams were created using BioRender (Biorender, RRID:SCR_018361). Graphs were generated in Graphpad Prism 10.6.1 (GraphPad Prism, Boston, MA; RRID:SCR_002798).

### Statistical analysis

Unless noted, no sex differences were seen and thus results were pooled for male and female mice. Data are expressed as means ± SEM and rounded to one decimal point. Reported percentages are rounded to one decimal point and thus may not add up to 100%. Significance was set at α<0.05. Statistical analysis was conducted using Graphpad Prism (10.6.1; GraphPad Software, San Diego, CA). Two-group comparisons were conducted using Student’s *t*-test.

## Results

### D1R

Reporter mouse studies indicate sparse, subregion-specific D1R expression in the hippocampus. In the CA1, D1R localization has been reported primarily in interneurons, whereas expression in pyramidal neurons remains controversial^[34, 43]^. In the CA3, D1R expression has been reported to be largely confined to interneurons, although localization in mossy fiber terminals has also been proposed^[34, 44]^. In the DG, D1R expression has been observed in granule cells as well as interneurons in the hilus and molecular layer^[34]^. Although these prior studies establish D1R expression across hippocampal subregions, they lack the ultrastructural resolution necessary to determine receptor localization within specific neuronal compartments that govern dopaminergic regulation of hippocampal function. To define the subcellular distribution of hippocampal D1R, we used immuno-electron microscopy to examine D1R localization in CA1 and CA3 SR, CA3 SLu, and DG hilus (Fig. 1) of male and female mice.

**Figure 1.**
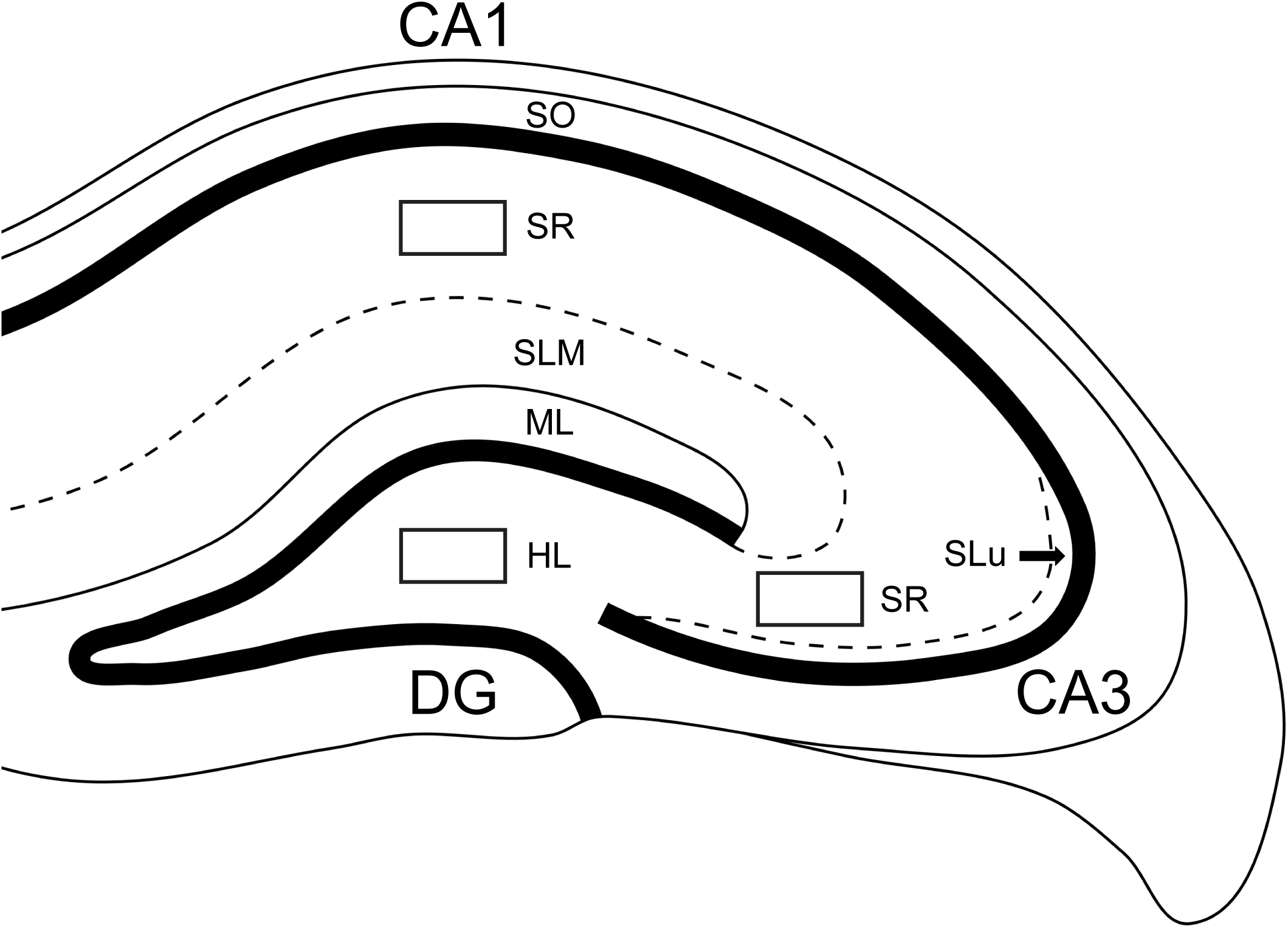
Sampling locations in the hippocampus. Images were sampled from 6050 µm2 (boxes) from the CA1, CA3, and dentate gyrus (DG). Abbreviations: hilus, HL; molecular layer, ML; stratum lacunosum-moleculare, SLM; stratum lucidum, SLu; stratum oriens, SO; stratum radiatum, SR. Mossy fibers were surveyed in the CA3 SLu.

### D1R in the CA1

Robust D1R immunoreactivity was observed in distinct profile patterns within the CA1 SR. Notably, D1R labeling was detected in dendritic spines (Fig. 2A, B), which were commonly observed (Fig. 2G), and typically formed synapses with terminal boutons (Fig. 2A, B) or *en passant* boutons (Fig. 2E). Among the 78 D1R labeled spines observed in the CA1, 60 (76.9%) received contacts from terminal boutons, 7 (9.0%) were innervated by *en passant* boutons, 2 (2.6%) formed synapses with unidentifiable profiles, and 9 (11.5%) showed no synaptic contact in the plane of section examined.

**Figure 2.**
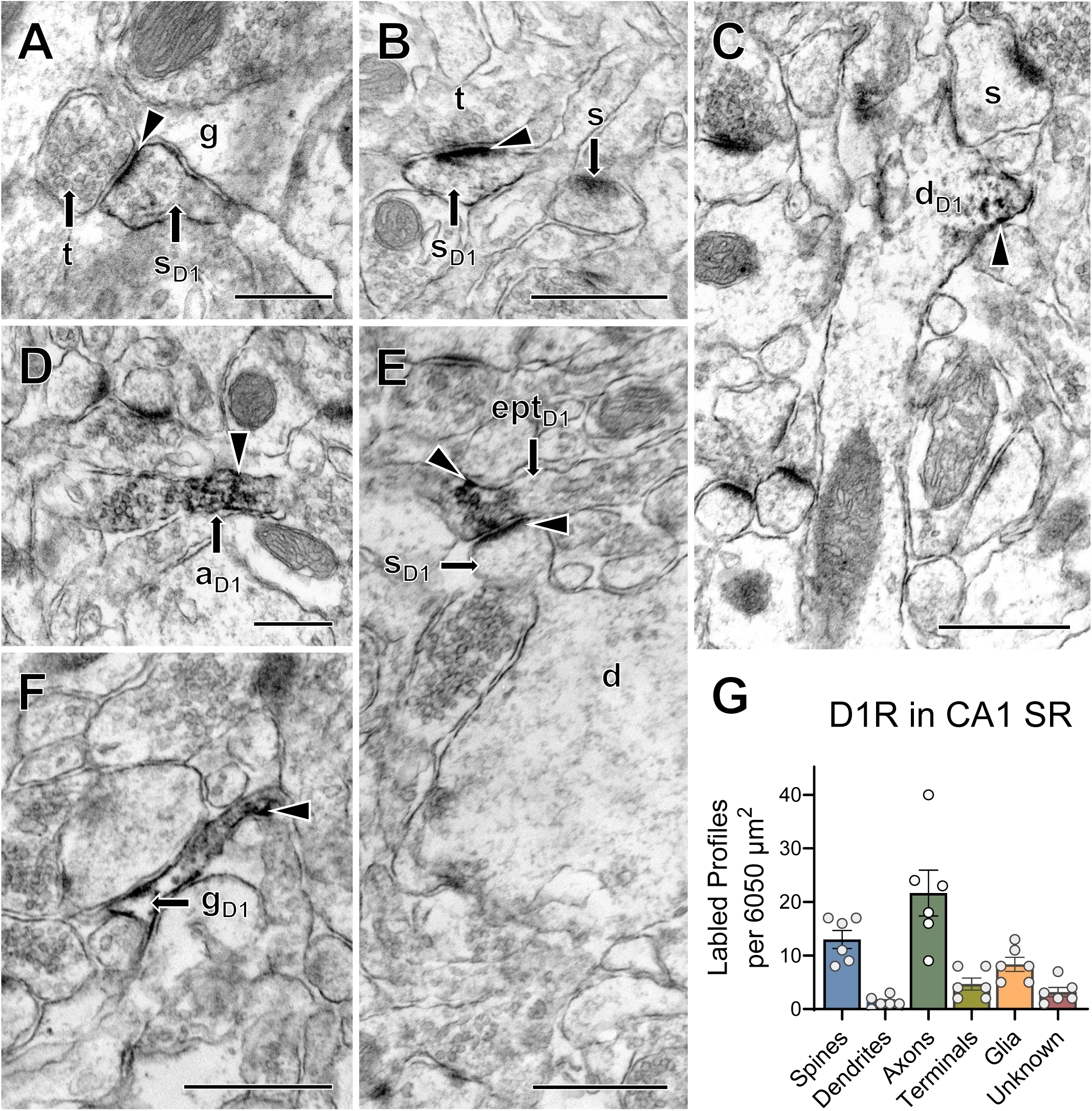
Ultrastructural distribution of dopamine 1 receptor labeled profiles in the CA1 SR. **A-F.** Representative images of D1R-labeled: **A,B)** dendritic spines (sd1) contacted by unlabeled terminal boutons (t), supported by unlabeled glia (g) or in proximity to unlabeled spine (s), **C)** pyramidal cell dendrite (dd1) with unlabeled spine (s), **D)** unmyelinated axon (ad1), **E)** en passant terminal (eptd1) forming asymmetric synapse with labeled dendritic spine (sd1), **F)** glia (gd1). Scale Bars = 400nm in A, D, 600nm in B, C, E, F. G. Total D1R-labeled profile prevalence. Data are expressed as mean +/-SEM, N = 3 female and 3 male mice.

Although D1R-labeled dendrites were observed in the CA1 (Fig. 2C), they were less numerous compared to labeled dendritic spines (Fig. 2G). Of the 8 D1R-labeled dendrites identified in the CA1, 7 (87.5%) displayed morphological characteristics of pyramidal neurons. Most D1R-labeled dendrites (62.5%) were not contacted by an identifiable profile in the plane of section analyzed, whereas two received contacts from *en passant* boutons and one from a terminal bouton.

D1R-labeled axons (Fig. 2D) were the most abundant labeled profile in the CA1 SR (Fig. 2G). Labeled axon terminals were present (Fig. 2E) but relatively rare (Fig. 2G). Of the 28 D1R-positive terminals, 17 (60.7%) formed asymmetric synapses, all of which targeted dendritic spines (**Table 2**). Symmetric synapses were less common and observed in 5 terminals (17.9%) (**Table 2**). These terminals contacted diverse dendritic targets: 3 (60%) innervated pyramidal dendrites, 1 (20%) contacted an interneuron dendrite, and 1 (20%) synapsed with a dendrite of unidentified cell-type. Five of the 28 terminals (17.9%) formed non-synaptic appositions (**Table 2**), most frequently contacting pyramidal dendrites (4 of 5; 80%), with 1 terminal contacting a dendritic spine (20%) (**Table 2**). Only 1 D1R-labeled axo-axonic terminal (3.6%) was observed in the CA1 SR (**Table 2**). Labeled terminals were typically bouton-type, although 7 *en-passant* boutons (25%) were also identified. D1R immunoreactivity was also detected in glial profiles (Fig. 2F), though at moderate prevalence (**Fig 2G**).

**Table 2:** D1R-labeled terminal contacts in the hippocampus.

|  | D1R-Labeled<br>Terminals | Spines |  |  | Dendrites |  |  | Unknown |  |  | Axo-Axo | No<br>Contact |
| --- | --- | --- | --- | --- | --- | --- | --- | --- | --- | --- | --- | --- |
|  |  | Asym | Sym | App | Asym | Sym | App | Asym | Sym | App |  |  |
| CA1 | 28 | 17 | 0 | 1 | 0 | 5 | 4 | 0 | 0 | 0 | 1 | 0 |
| CA3 | 27 | 15 | 0 | 3 | 1 | 1 | 1 | 1 | 1 | 0 | 2 | 1 |
| DG | 10 | 5 | 0 | 0 | 0 | 3 | 2 | 0 | 0 | 0 | 0 | 0 |

### D1R in the CA3

D1R-labeled profile patterns in the CA3 SR were similar to those observed in the CA1. D1R labeling was detected in dendritic spines (Fig. 3A), indicating expression by CA3 pyramidal neurons (Fig. 3H). Of the 48 D1R-labeled spines observed, 41 (85.4%) received innervation from terminal boutons, 5 (10.4%) received contacts from *en passant* boutons, and 2 (4.2%) showed no synaptic contact in the examined plane of section. D1R-labeled dendritic shafts were rarely observed in the CA3. Only one labeled dendrite was identified across six sampled mice (Fig. 3H); this profile exhibited morphological characteristics of a pyramidal neuron and showed no observable synaptic contact in the observed section. D1R labeling was also detected in axons (Fig. 3B, C), which represented the most abundantly labeled profile in the CA3 SR (Fig. 3H). Females tended to exhibit a higher prevalence of labeled axons than males, although this difference did not reach statistical significance (**Supplementary Fig. 1**).

**Figure 3.**
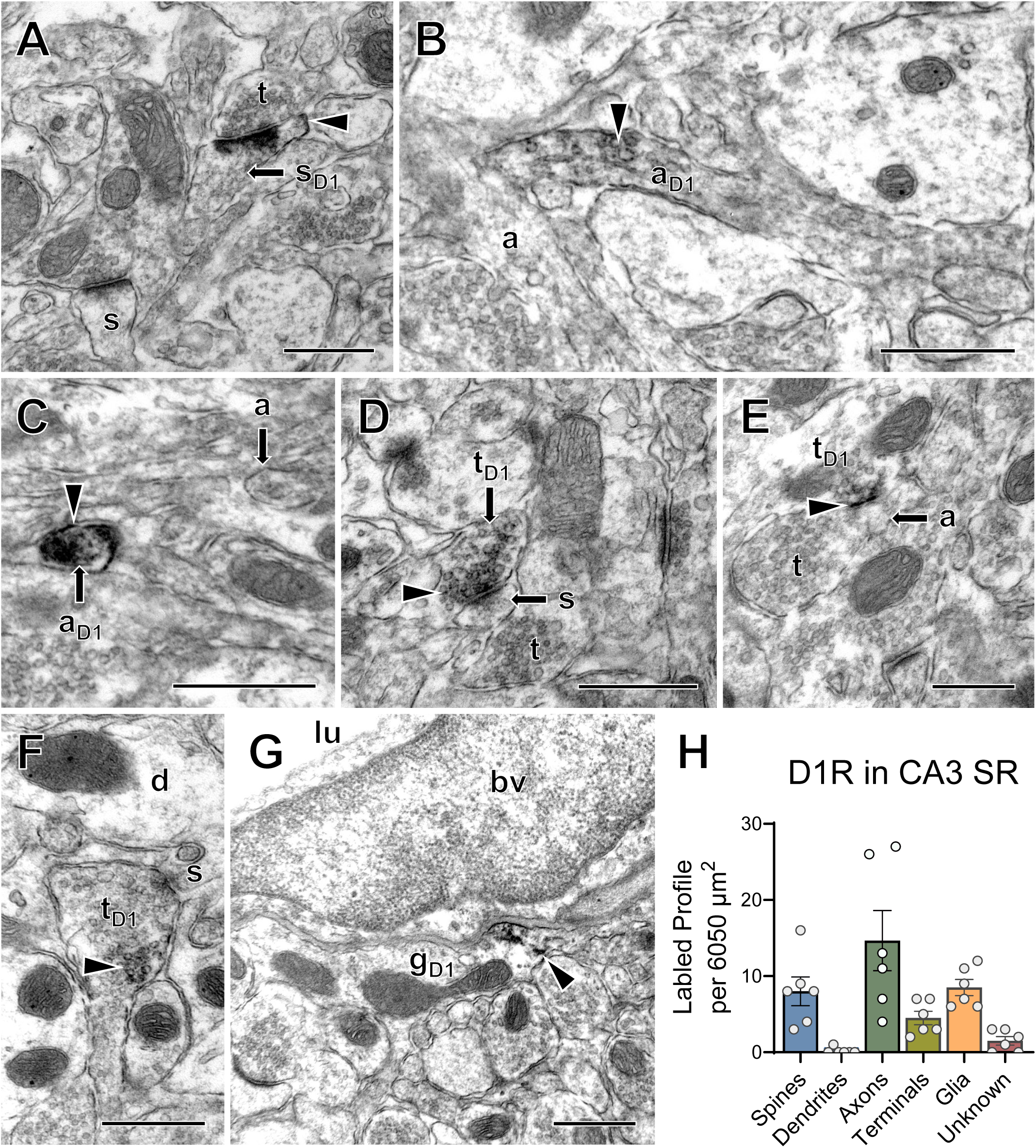
Ultrastructural distribution of dopamine 1 receptor labeled profiles in the CA3 SR. **A-G.** Representative images of D1R-labeled: **A)** dendritic spine (sd1) contacted by unlabeled terminal bouton (t) in proximity to unlabeled spine (s), **B,C)** unmyelinated axon (ad1), **D)** axon terminal bouton (td1) forming asymmetric synapse with unlabeled spine (s), **E)** axon terminal bouton (td1) forming axo-axonic synapse with unlabeled axon (a) immediately preceding an unlabeled terminal bouton (t), **F)** axon terminal bouton (td1) forming apposition with dendritic spine (s), and **G)** glia (gd1) apposing blood vessel with visible lumen (lu). Scale Bars = 400nm in E, 600 nm in A, B, C, D, F, G. **H.** Total D1R-labeled profile prevalence. Data are expressed as mean +/- SEM, N = 3 female and 3 male mice.

D1R-labeled axon terminals were present but relatively uncommon in the CA3 SR (Fig. 3H). Of the 26 labeled terminals analyzed, 17 (65.4%) formed asymmetric synapses (Fig. 3D; **Table 2**). These terminals primarily targeted dendritic spines (15 of 17; 88.2%), with the remainder contacting a pyramidal dendrite (1 of 17; 5.9%) or an unidentifiable profile (1 of 17; 5.9%) (**Table 2**). Symmetric synapses were less frequent and were observed in only 2 terminals (7.7%) (Fig. 3F; **Table 2**). Five terminals (19.2%) formed appositions, contacting dendritic spines (3 of 5; 60%), dendrites of unidentifiable cell type (1 of 5; 20%), or unidentifiable profiles (1 of 5; 20%) (**Table 2**). Four terminals (15.4%) formed *en passant* synapses. Two axo-axonic D1R-labeled terminals were observed (7.7%; Fig. 3E; **Table 2**), both in female animals. Glial labeling was detected in the CA3 SR (Fig. 3G) but occurred at a lower prevalence (Fig. 3H).

To assess previous reports of D1R expression in mossy fiber synapses, 50 mossy fiber terminals per animal were examined in the SLu. Only 1.7% of mossy fiber terminals exhibited D1R labeling (data not shown), suggesting that D1R contribution to mossy fiber synaptic plasticity is limited.

### D1R in DG

D1R labeling patterns in the DG hilus differed from those observed in the CA1 and CA3 SR. Labeling was detected predominantly in axons (Fig. 4B), and glial profiles (Fig. 4C). Interestingly, females tended to exhibit a higher prevalence of D1R-labeled axons than males, although this difference did not reach statistical significance (**Supplementary Figure 1**). D1R labeling in dendritic spines and dendritic shafts was rare in the hilus (Fig. 4D) and only a single labeled dendrite was identified, which displayed morphological characteristics of an interneuron. D1R-labeled axon terminals were infrequent in the hilus (**Table 2**). Of the 10 labeled terminals analyzed, 5 (50%) formed asymmetric synapses, all of which targeted dendritic spines (**Table 2**). Symmetric synapses were identified in 3 terminals (30%), each contacting dendrites with morphological characteristics of mossy cells (**Table 2**). Two terminals (20%) formed nonsynaptic appositions, contacting either a mossy cell dendrite or a dendrite of unidentifiable cell type. One terminal (10%) formed an *en passant* synapse. A single mossy fiber terminal with D1R labeling was observed in the DG (Fig. 4A), indicating rare labeling of mossy fiber projections. D1R immunoreactivity was also observed in glial profiles (Fig. 4C) but was infrequent (Fig. 4D).

**Figure 4.**
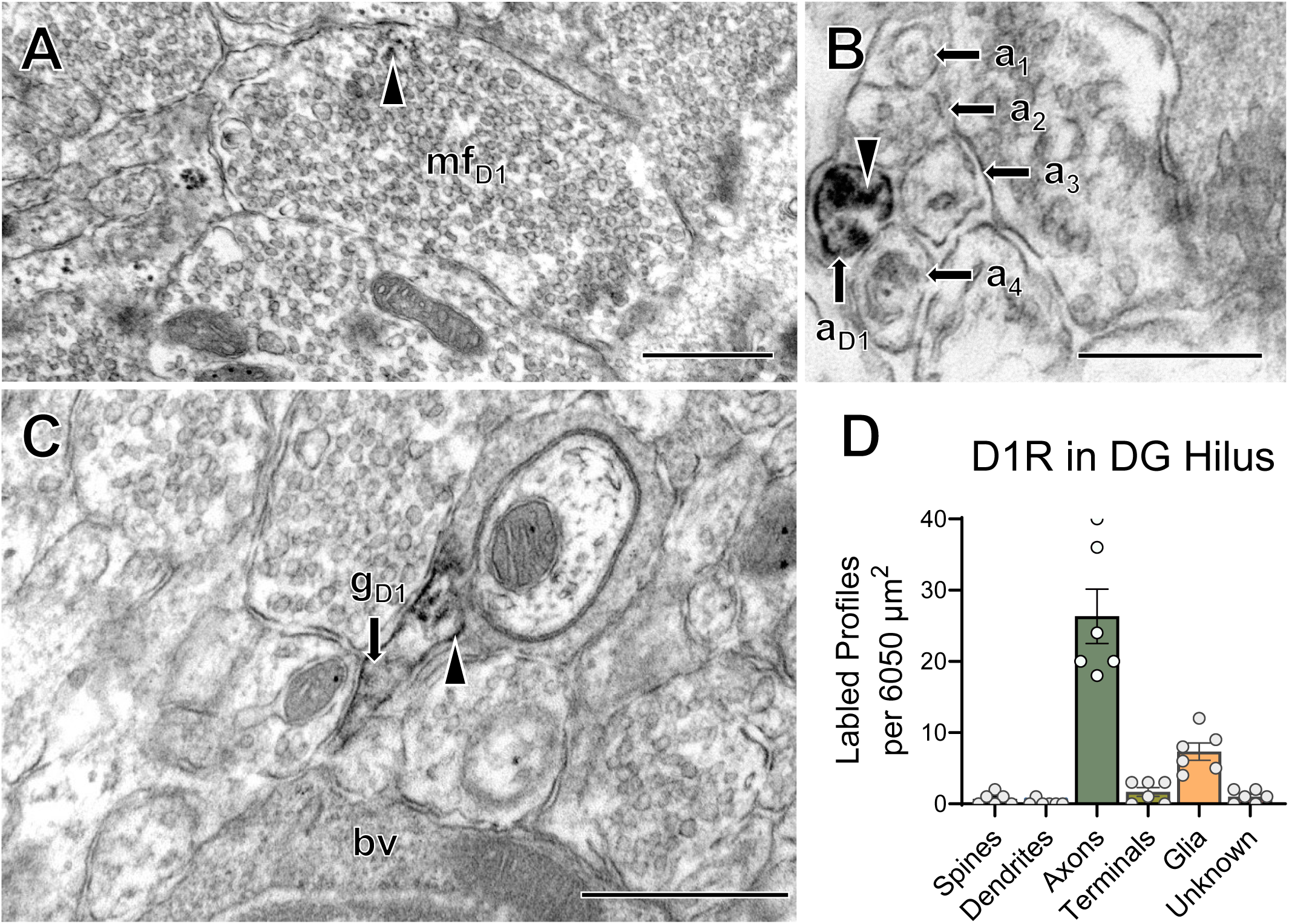
Ultrastructural distribution of dopamine 1 receptor labeled profiles in the DG HL. **A-G.** Representative images of D1R-labeled: **A)** mossy fiber (mfd1), **B)** unmyelinated axon (ad1) in proximity to unlabeled axons (a1, a2), and **C)** glia (gd1) in proximity to blood vessel (bv). Scale Bars = 400nm in B, 600nm in A, C. **D.** Total D1R-labeled profile prevalence. Data are expressed as mean +/- SEM, N = 3 female and 3 male mice.

### D2R

Previous reporter mouse-based studies suggest a more restricted pattern of D2R expression in the hippocampus. In the CA1 and CA3, D2R labeling has been reported primarily in interneurons ^[35]^, whereas expression in principal neurons remains uncertain. In the DG, D2R expression has been described in hilar mossy cells^[34, 35]^ and has not been widely detected in other cell populations. To determine the subcellular distribution of hippocampal D2R, we used immunoelectron microscopy to examine D2R localization in the CA1 and CA3 SR, CA3 SLu, and the DG hilus of male and female mice (Fig. 1).

### D2R in the CA1

In the CA1 SR, D2R labeling was detected in dendritic spines (Fig. 5A, D), which were abundant (Fig. 5H). Females tended to have a higher prevalence of D2R-labeled spines than males, although this difference did not reach statistical significance (**Supplemental Fig. 1**). Of the 89 D2R-labeled spines analyzed, 76 (85.4%) received contacts from terminal boutons, 7 (7.9%) were contacted by *en passant* boutons, and 6 (6.7%) showed no synaptic contact in the plane of section examined. D2R-labeled dendritic shafts also were observed in the CA1 (Fig. 5B, E) but were relatively infrequent (Fig. 5H). Of the 7 labeled dendrites identified, 4 (57.1%) showed no synaptic contact. Most of these uncontacted dendrites (3 out of 4; 75%) were adjacent to unlabeled axons (Fig. 5B). Two dendrites (28.6%) received innervation from *en passant* boutons and 1 (14.3%) from a terminal bouton. Four labeled dendrites (57.1%) exhibited morphological characteristics of pyramidal neurons, one (14.3%) corresponded to an interneuron dendrite, and two (28.6%) were classified as dendrites of unidentifiable cell-type.

**Figure 5.**
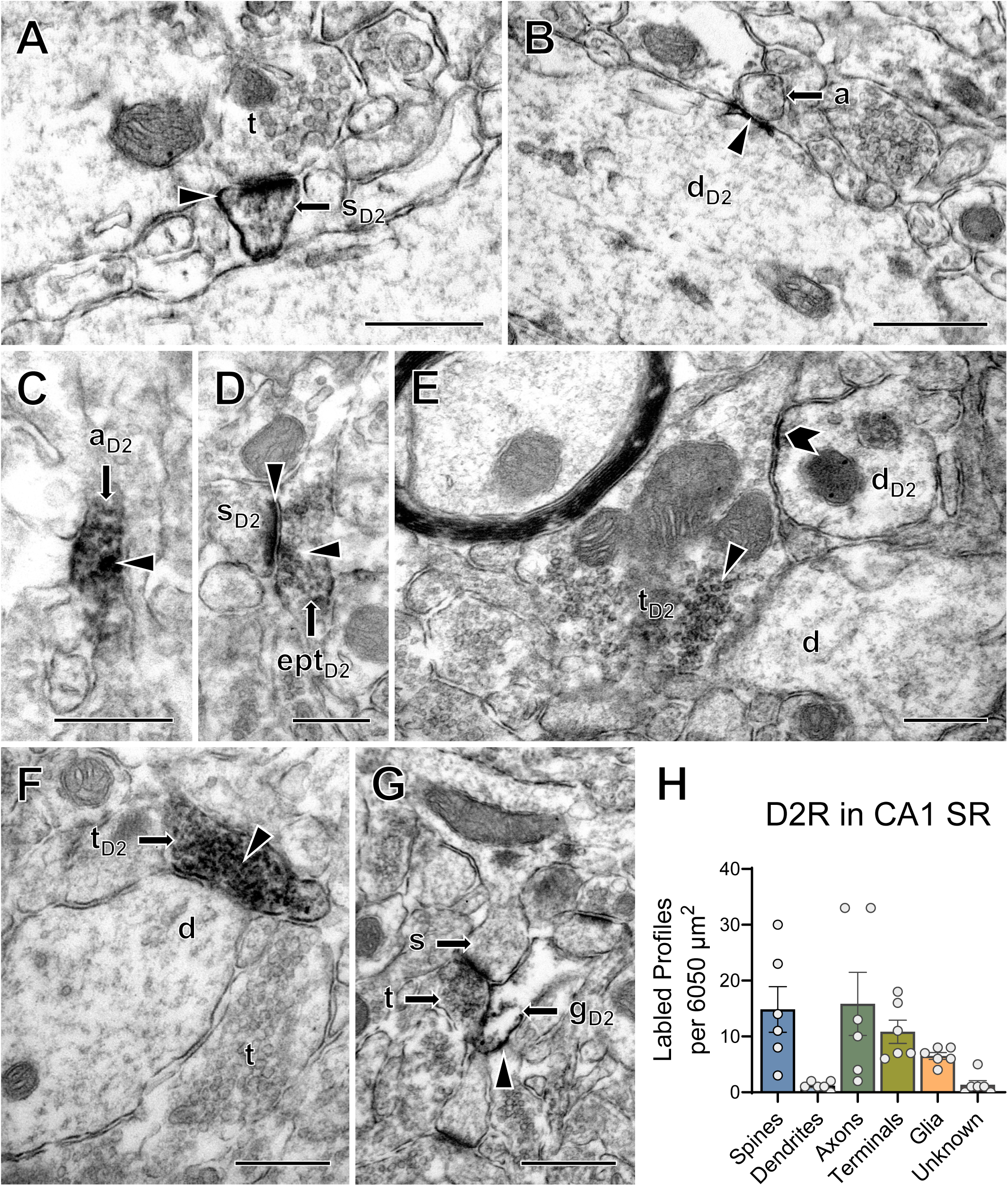
Ultrastructural distribution of dopamine 2 receptor labeled profiles in the CA1 SR. **A-G.** Representative images of D2R-labeled: A) dendritic spine (sd2) contacted by an unlabeled terminal bouton (t), **B)** pyramidal cell dendrite (dd2) in proximity to unlabeled axon (a), **C)** unmyelinated axon (ad2), **D)** en-passant axon terminal (eptd2) forming an asymmetric synapse with a labeled dendritic spine (sd2), **E)** axon terminal bouton (td2) forming an apposition with an unlabeled dendrite (unknown cell type) (d) in proximity to labeled dendrite of unknown cell type (dd2, chevron), F) axon terminal bouton (td2) forming a symmetric synapse with an unlabeled interneuron dendrite (d), and G) Glia (gd2) supporting a synapse between unlabeled axon terminal bouton (t) and dendritic spine (s). Scale Bars = 400nm in A, C, D, E, F, G, 600nm in B. H. Total D2R-labeled profile prevalence. Data are expressed as mean +/- SEM, N = 3 female and 3 male

D2R labeling also was detected in axons (Fig. 5C) and were frequently observed in the CA1 SR (Fig. 5H). D2R-labeled axon terminals were abundant and formed asymmetric synapses (Fig. 5D), appositions (Fig. 5E), and symmetric synapses (Fig. 5F), occasionally in *en passant* configurations (Fig. 5D). Of the 65 labeled terminals with identifiable synapses, 29 (46.1%) formed symmetric synapses (**Table 3**). These terminals primarily contacted pyramidal neuron dendrites (21 of 29; 75.9%), with fewer targeting dendrites of unidentifiable cell type (6 of 29; 20.7%) or interneuron dendrites (1 of 29; 3.4%). Sixteen terminals (24.6%) formed asymmetric synapses (**Table 3**), almost exclusively onto dendritic spines (15 of 16; 93.8%), with one contacting an interneuron dendrite (1 of 16; 6.3%). In addition, 12 labeled terminals (18.5%) formed appositions (**Table 3**), contacting pyramidal dendrites (8 of 12; 66.7%), dendrites of unidentifiable cell type (2 of 12; 16.7%), an interneuron dendrite (1 of 12; 8.3%), or a dendritic spine (1 of 12; 8.3%). Axo-axonic synapses were rare, with only 2 observed (3.1%) (**Table 3**). *En passant* synapses were also uncommon (5 of 65; 6.2%). D2R immunoreactivity was detected in glial profiles (Fig. 5G) but occurred at low prevalence (Fig. 5H).

**Table 3:**
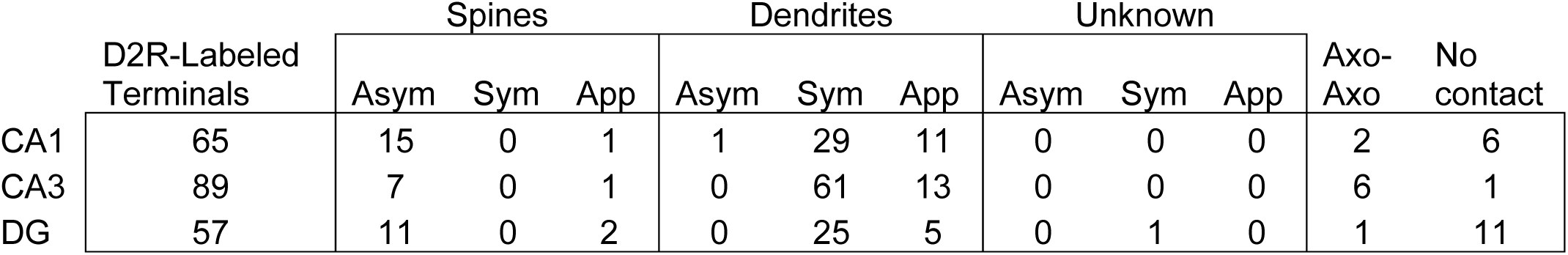
D2R-labeled terminal contacts in the hippocampus.

|  | D2R-Labeled<br>Terminals | Spines |  |  | Dendrites |  |  | Unknown |  |  | Axo-Axo | No<br>contact |
| --- | --- | --- | --- | --- | --- | --- | --- | --- | --- | --- | --- | --- |
|  |  | Asym | Sym | App | Asym | Sym | App | Asym | Sym | App |  |  |
| CA1 | 65 | 15 | 0 | 1 | 1 | 29 | 11 | 0 | 0 | 0 | 2 | 6 |
| CA3 | 89 | 7 | 0 | 1 | 0 | 61 | 13 | 0 | 0 | 0 | 6 | 1 |
| DG | 57 | 11 | 0 | 2 | 0 | 25 | 5 | 0 | 1 | 0 | 1 | 11 |

### D2R in CA3

In the CA3 SR, D2R labeling was detected in dendritic spines (Fig. 6A), but not in dendritic shafts (Fig. 6H). Labeled spines were abundant (Fig. 6H), indicating postsynaptic expression in CA3 pyramidal neurons. Of the 61 labeled spines analyzed, 44 (72.1%) received contacts from terminal boutons (Fig. 6A), 11 (18%) were contacted by *en passant* boutons, 1 (1.6%) contacted an unidentifiable profile, and 5 (8.2%) showed no synaptic contact in the plane of section examined.

**Figure 6.**
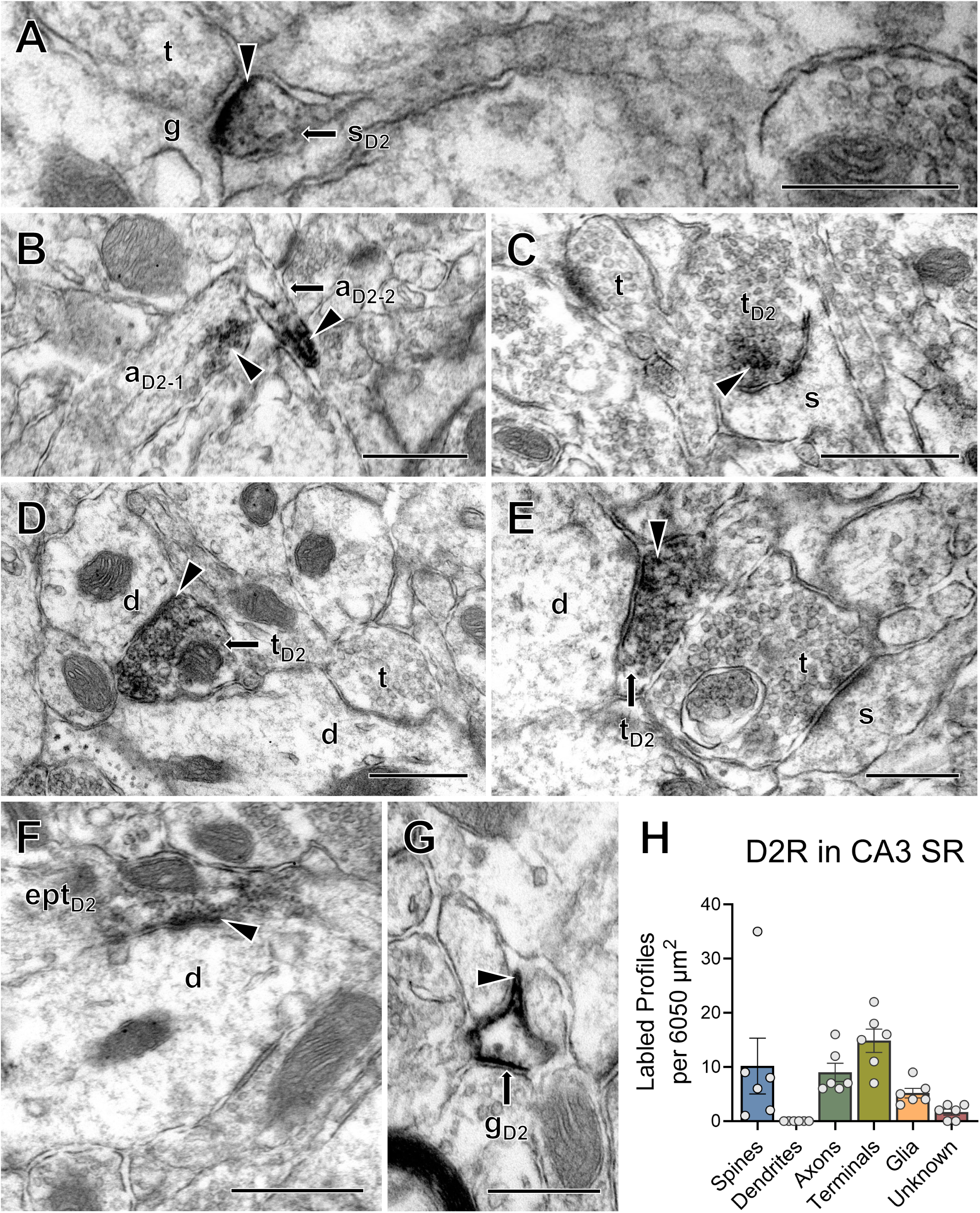
Ultrastructural distribution of dopamine 2 receptor labeled profiles in the CA3 SR. **A-G.** Representative images of D2R-labeled: **A)** dendritic spine (sd2) contacted by an unlabeled terminal bouton (t), B) unmyelinated axons (ad2-1, ad2-2) C) axon terminal bouton (td2) forming an asymmetric synapse with an unlabeled spine (s), **D,E)** axon terminal boutons (td2) forming a symmetric synapse with unlabeled pyramidal cell dendrites (d) next to unlabeled terminals (t) and their targets (d,s), **F)** en passant axon terminal (eptd2) forming symmetric synapse with unlabeled pyramidal cell dendrite (d), and **G)** glia (gd2). Scale Bars = 400nm in A, E, F, G, 600nm in B, C, D. **H.** Total D2R-labeled profile prevalence. Data are expressed as mean +/- SEM, N = 3 female and 3 male mice.

D2R labeling was also observed in axons (Fig. 6B), which were relatively common in the CA3 SR (Fig. 6H). Axon terminals represented the most prevalent labeled profile and formed asymmetric synapses (Fig. 6C), symmetric synapses (**Fig. 6D-F**), and appositions. Of the 89 labeled terminals analyzed, 61 (68.5%) formed symmetric synapses (**Table 3**). These terminals primarily contacted pyramidal neuron dendrites (50 of 61; 82%), with fewer targeting dendrites of unidentifiable cell-type (10 of 61; 16.4%), or interneuron dendrites (1 of 61; 1.6%). Asymmetric synapses were less frequent and were observed in 7 terminals (7.9%) (**Table 3**), all of which contacted dendritic spines. These terminals were predominantly observed in female animals (data not shown). Fourteen terminals (15.7%) formed appositions (**Table 3**), contacting pyramidal neuron dendrites (10 of 14; 71.4%), dendrites of unidentifiable cell type (3 of 14; 21.4%), or, in one case, a dendritic spine (1 of 14; 7.1%). *En passant* synapses were rare, occurring in only 6 terminals (6.7%). Axo-axonic synapses were also uncommon, with 6 observed (6.7%), all in female mice (**Table 3**). D2R immunoreactivity was detected in glial profiles (Fig. 6G), though at low prevalence (Fig. 6H).

To assess D2R expression in mossy fiber terminals, 50 mossy fibers per animal were examined in the SLu. D2R labeling was detected only in male mossy fibers and in a small proportion overall, with 4% of mossy fibers exhibiting labeling (data not shown).

### D2R in DG

In the DG central hilus, D2R labeling exhibited a pattern more similar to that observed in the CA1 and CA3 SR than hilar D1R labeling. D2R immunoreactivity was detected in dendritic spines (Fig. 7A), consistent with postsynaptic expression in hilar mossy cells, although labeled spines were relatively rare (Fig. 7G). Of the 21 labeled spines analyzed, 19 (90.5%) received contacts from terminal boutons, 1 (4.8%) was innervated by an *en passant* bouton, and 1 (4.8%) showed no synaptic contact in the plane of section examined. D2R-labeled dendritic shafts also were observed (Fig. 7B) but occurred at low prevalence (Fig. 7G). These dendrites received contacts from a variety of profiles: 4 (36.4%) formed synapses with mossy fiber terminals, 3 (27.3%) with terminal boutons, 1 (9.1%) with an *en passant* bouton, and 1 (9.1%) with an unidentifiable profile, whereas 2 (18.2%) showed no observable synapse in the sampled plane. Due to the dense CA3 dendritic projections into the central hilus, definitive identification of dendritic cell type was often not possible. Only one labeled dendrite (9.1%) was identified as belonging to an interneuron, whereas the remaining profiles were classified as dendrites of unidentifiable cell type.

**Figure 7.**
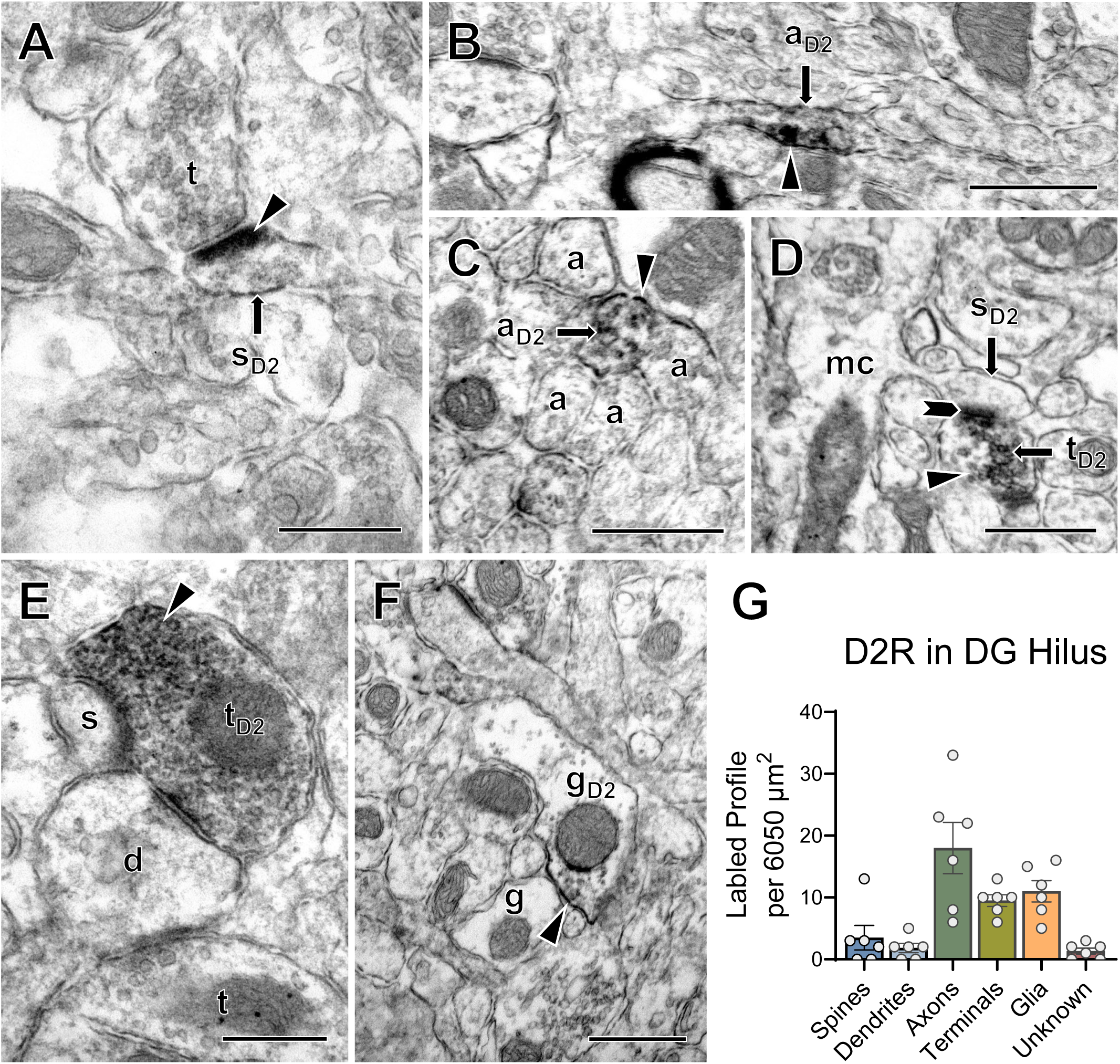
Ultrastructural distribution of dopamine 2 receptor labeled profiles in the DG HL. **A-F.** Representative images of D2R-labeled: **A)** dendritic spine (sd2) contacted by an unlabeled terminal bouton (t), **B,C)** unmyelinated axon (ad2), **D)** axon terminal (td2) forming asymmetric synapse with labeled mossy cell spine (sd2), **E)** axon terminal bouton (td2) forming symmetric synapse with unlabeled dendrite of unknown cell type (d), and apposition with unlabeled spine (s), and **F)** glia (gd2). Scale Bars = 400nm in A, E, 600nm in B, C, D, F. **G.** Total D2R-labeled profile prevalence. Data are expressed as mean +/- SEM, N = 3 female and 3 male mice.

Axons represented the most commonly observed labeled profile in the hilus (Fig. 7B, C, G). Labeled axon terminals formed asymmetric synapses (Fig. 7D), symmetric synapses (Fig. 7E), and appositions (Fig. 7G). Of the 57 labeled terminals analyzed, 26 (45.6%) formed symmetric synapses (**Table 3**). These terminals contacted interneuron dendrites (11 of 26; 42.3%) mossy cell dendrites (2 of 26; 7.7%), or dendrites of unidentifiable cell type (12 of 26; 46.2%). Asymmetric synapses were less frequent and were observed in 11 terminals (19.3%) (**Table 3**), all of which targeted dendritic spines. Seven terminals (12.3%) formed appositions (**Table 3**), contacting spines (2 of 7; 28.6%), mossy cell dendrite (1 of 7; 14.3%), or dendrites of unidentifiable cell type (4 of 7; 57.1%). Only one axo-axonic terminal (1.8%) was observed. Most terminals were bouton-type endings, with only 3 (5.3%) forming *en passant* synapses. D2R immunoreactivity also was detected in glial profiles (Fig. 7F), which were relatively common in the hilus (Fig. 7G).

## Discussion

This study describes the subcellular distribution of D1R- and D2R-containing profiles across hippocampal subregions and identifies distinct cellular and synaptic localization patterns for each receptor. In CA1 and CA3 SR, both D1R and D2R immunoreactivity was prominently localized to pyramidal neuron compartments, including dendritic spines and unmyelinated axons, whereas receptor labeling in axon terminals and glia occurred at lower prevalence, Notably, D1R-and D2R-containing terminals formed different classes of synapses, with D1R associated primarily with excitatory-type contacts on dendritic spines and D2R associated predominantly with inhibitory-type contacts on pyramidal neuron dendritic shafts. In the DG hilus, D1R labeling was sparse and largely confined to axons and glia, whereas D2R labeling was more broadly distributed, appearing on mossy cell spines and inhibitory terminals. Together, these findings indicate that dopamine receptors are strategically positioned to modulate excitatory and inhibitory signaling within hippocampal circuits and provide a structural substrate for bidirectional dopaminergic regulation of synaptic plasticity and neuronal activity.

### Postsynaptic dopamine receptors mediate synaptic plasticity

In CA1 and CA3 SR, both DR1- and DR2-immunoreactivities were concentrated in dendritic spines of pyramidal neurons, which receive predominantly excitatory glutamatergic input. In CA1 SR, these inputs arise mostly from Schaffer collateral projections originating in CA3^[45]^, whereas in CA3 SR excitatory drive is provided largely by associational and commissural recurrent collateral pathways^[46]^. The prominent localization of D1R and D2R within pyramidal neuron dendritic spines suggest that dopaminergic signaling can directly modulate glutamatergic transmission and synaptic plasticity at these synapses. Activation of postsynaptic D1Rs enhances neuronal excitability, strengthens synaptic transmission, and facilitates long-term potentiation (LTP) through multiple mechanisms, including increased NMDA and AMPA receptor currents^[11, 17, 47]^, enhanced temporal summation during high-frequency input^[16]^, promotion of receptor trafficking to the plasma membrane^[25, 26]^, and reduced inhibitory GABA_A_ receptor conductance^[18]^. In contrast, postsynaptic D2R activation is generally associated with long-term depression (LTD), mediated in part by inhibition of cAMP signaling^[10]^, reductions in AMPA and NMDA receptor currents ^[48–51]^, and decreased membrane insertion of AMPA receptors^[26]^. The labeling of both receptor subtypes within pyramidal neuron spines therefore provides a molecular basis through which dopaminergic signaling may bias excitatory synapses toward potentiation or depression depending on receptor engagement and local dopamine availability.

In the DG hilus, D2R labeling was detected in mossy cell dendritic spines receiving excitatory input that likely arises from granule cell mossy fibers and CA3 pyramidal neurons^[52]^. This postsynaptic localization suggests that dopamine may directly regulate mossy cell excitability and the feedback circuits that control granule cell activity. As D2Rs typically reduce neuronal excitability though inhibition of cAMP-dependent signaling^[10]^, activation of these receptors on mossy cells may dampen excitatory drive within the hilus.

### Presynaptic dopamine receptors regulate excitatory and inhibitory transmission

D1R- and D2R-labeling in axons and presynaptic terminals indicates that dopamine can also modulate hippocampal signaling through presynaptic mechanisms at specific inputs. In CA1 and CA3, labeling for both receptors were detected on excitatory-type terminals forming synapses onto pyramidal neuron dendritic spines. These terminals most likely project from CA3 pyramidal neurons via the Schaffer collateral pathway in CA1, or from associational and commissural recurrent collateral pathways within CA3^[53]^. Dopaminergic modulation of these terminals can bidirectionally regulate glutamate release and thereby influence hippocampal synaptic signaling and plasticity. Activation of presynaptic D1R enhances glutamate release^[8]^, whereas D2R activation reduces glutamate release^[54]^. Thus, in addition to postsynaptic mechanisms of plasticity, dopaminergic signaling may regulate pyramidal neuron activity by modulating excitatory afferent drive.

Dopamine may also influence inhibitory signaling within the hippocampus. The activity of pyramidal neurons and mossy cells is shaped by diverse populations of GABAergic interneurons as well as inhibitory inputs from extrahippocampal regions, including the medial septum^[53]^. Our results demonstrate frequent association of D2R labeling with terminals forming symmetric synapses throughout the hippocampus. Though reports predominantly suggest that activation of presynaptic D2R reduces GABA release^[55]^, this effect may be cell-type specific^[14]^. The localization of D2R observed here suggests that dopaminergic signaling may regulate inhibitory transmission within hippocampal circuits, although further studies are required to confirm this effect.

### D1- and D2-receptor signaling in hippocampal learning and memory

Hippocampal dopamine plays a critical role in learning and memory processes, including object location memory, spatial navigation, fear conditioning, contextual memory linking, memory updating, and encoding of novel information^[20, 22, 28, 29, 56]^. In the dorsal hippocampus, dopamine arises predominantly from locus coeruleus (LC) noradrenergic afferents rather than ventral tegmental area (VTA) projections^[28]^. LC neurons exhibit distinct firing patterns, including low-frequency tonic activity that maintains baseline extracellular catecholamine levels and high-frequency phasic bursts that generate transient elevations in dopamine^[57, 58]^. Given that D2R have a higher affinity for dopamine than D1R, tonic dopamine release is thought to preferentially engage D2Rs, whereas robust D1R activation requires the higher concentrations associated with phasic release^[59, 60]^. Consistent with this framework, dopamine-dependent learning and memory processes are strongly associated with phasic LC activity^[58, 61]^, and consequent activation of hippocampal D1R^[20, 22, 28]^.

Our results provide a potential circuit mechanism linking phasic LC firing to learning and memory behavior. The receptor distributions observed here suggest that tonic LC activity may preferentially modulate inhibitory inputs and reduce the excitability of pyramidal neurons and mossy cells through D2R activation. In contrast, phasic dopaminergic signaling would additionally recruit lower-affinity D1R, enhancing excitatory transmission and facilitating activity-dependent plasticity at select pyramidal neurons. Localization of D1R within pyramidal neuron spines, together with selective expression in excitatory terminals, suggests that transient dopamine elevations associated with phasic LC firing could bias active synapses toward potentiation during salient events, thereby facilitating memory encoding.

### Sex differences in hippocampal dopamine receptor expression

Although not statistically significant, we observed a trend toward higher numbers of D1R-and D2R-labeled profiles in females across most hippocampal regions examined, with the exception of D2R labeling in the DG. These observations contrast with reporter mouse-based findings suggesting reduced hippocampal D2R expression in females^[62]^, highlighting the importance of direct immunohistological studies. Differences in receptor expression may contribute to previously reported sex-dependent behavioral phenotypes. In spatial navigation paradigms, male rodents often employ different search strategies and may outperform females^[63–65]^. Similarly, males may exhibit stronger fear memory in conditioning paradigms, although this difference may partly reflect variation in fear behavior expression^[66]^. Though such behaviors are known to involve hippocampal dopamine signaling^[28, 56]^, prior work has mainly focused on sex hormones as the primary explanation for these differences^[67]^. Our results suggest that sex-dependent dopamine receptor distribution may represent an additional anatomical basis contributing to these behavioral effects. Although the functional significance of these observations remains to be determined, increased receptor availability could influence dopaminergic modulation of synaptic plasticity and circuit activity. These findings highlight sex-dependent dopamine receptor distribution as a potential factor underlying variability in dopamine-driven learning and reward-related behaviors and point to an important direction for future investigation.

In summary, this study identifies key similarities and differences in the subcellular localization of D1R and D2R across the CA1, CA3, and DG. The presence of both receptors on pyramidal neuron dendritic spines receiving excitatory inputs suggests that dopamine bidirectionally modulates excitatory signaling within hippocampal circuits. In addition, the selective localization of D2R on inhibitory terminals indicates that dopamine may also regulate inhibitory tone, thereby gating hippocampal function.

## Acknowledgements

Supported by NIH grants R01 GM130722 (JP), R01 HL136520 (TAM) and T32 DA039080 (CS). R.S. was supported by the WCM Summer Program for Science Teachers. The Hitachi HT7800 transmission microscope was obtained from HNIH Office of Research Infrastructure Program S10 Instrumentation Grant # 1S10OD026974-01A. We thank the Neuroanatomy EM Core at Weill Cornell for technical assistance.

## Data availability

Data is available upon request

## Ethics Approval

All ethics and integrity policies have been upheld by all listed authors. All material was personally created.

