## Supplemental Figure 1 for "Subcellular Localization of Dopamine D1 and D2 Receptors in the Mouse Hippocampus"

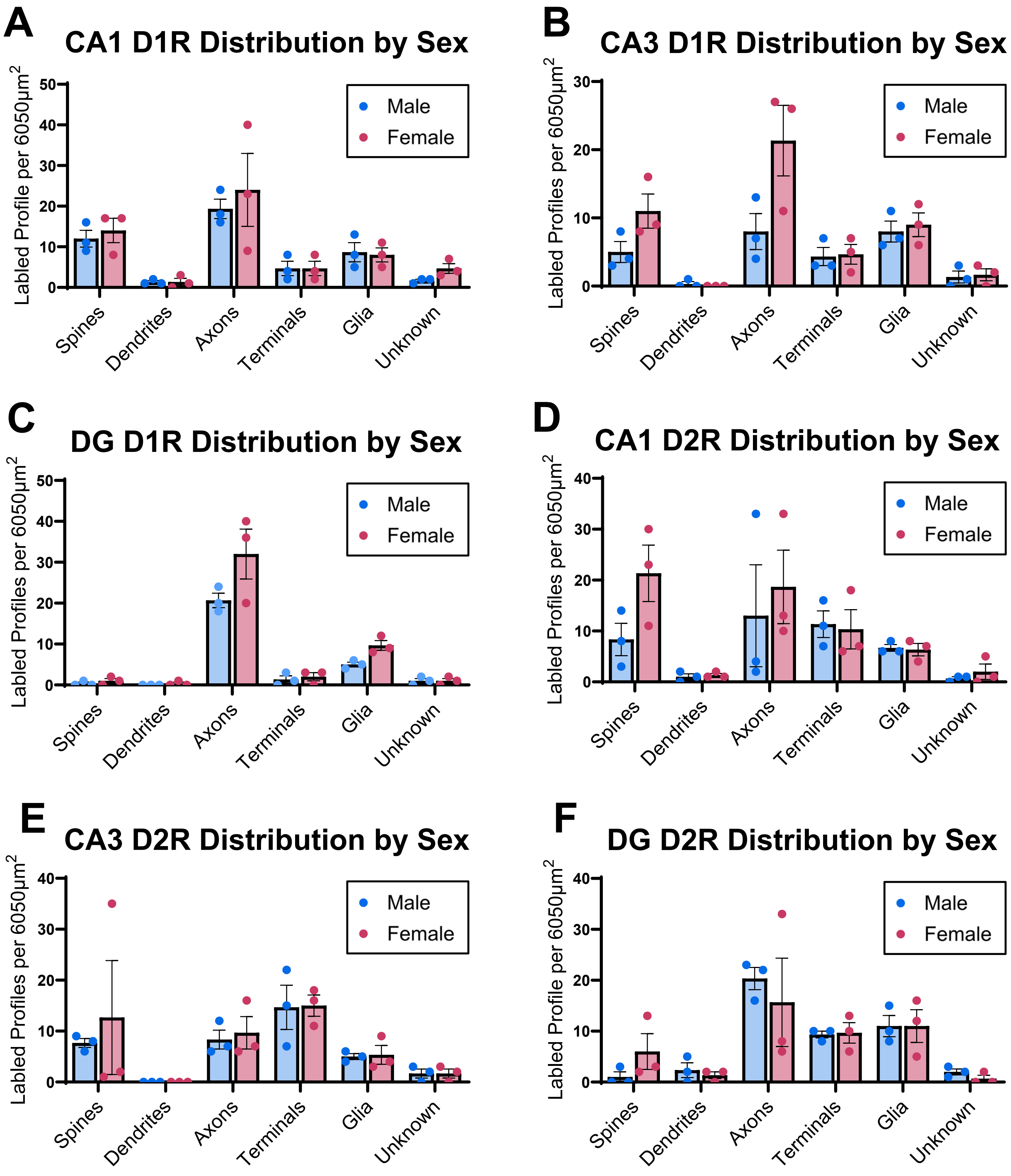


**Supplementary Figure 1. Sex differences of DAR labeled profiles in the hippocampus. A-C.** Sex differences in the prevalence of D1R labeled profiles in the **A)** CA1 SR, **B)** CA3 SR, and **C)** DG HL. **D-F.** Sex differences in the prevalence of D2R labeled profiles in the **D)** CA1 SR, **E)** CA3 SR, and **F)** DG HL.
